# Increased gene dosage and mRNA expression from chromosomal duplications in *C. elegans*

**DOI:** 10.1101/2022.03.22.485368

**Authors:** Bhavana Ragipani, Sarah Elizabeth Albritton, Ana Karina Morao, Diogo Mesquita, Maxwell Kramer, Sevinç Ercan

## Abstract

Isolation of copy number variations and chromosomal duplications at high frequency in the laboratory suggested that *Caenorhabditis elegans* tolerates increased gene dosage. Here we addressed if a general dosage compensation mechanism acts at the level of mRNA expression in *C. elegans*. We characterized gene dosage and mRNA expression in three chromosomal duplications and a fosmid integration strain using DNA-seq and mRNA-seq. Our results show that on average, increased gene dosage leads to increased mRNA expression, pointing to a lack of genome-wide dosage compensation. Different genes within the same chromosomal duplication show variable levels of mRNA increase, suggesting feedback regulation of individual genes. Somatic dosage compensation and germline repression reduce the level of mRNA increase from X chromosomal duplications. Together, our results show a lack of genome-wide dosage compensation mechanism acting at the mRNA level in *C. elegans* and highlight the role of epigenetic and individual gene regulation contributing to the varied consequences of increased gene dosage.

## INTRODUCTION

Chromosomal aneuploidies or copy number variations are associated with a wide range of phenotypes in many organisms (Tang and Amon 2013; Durrbaum and Storchova 2016). To understand the effect of gene dosage on gene expression, a series of studies compared DNA copy number and mRNA levels in aneuploid cells and animals (Kojima and Cimini 2019). In plants and mammalian cells, partial and full aneuploidies showed complex responses to alterations in gene dosage, with secondary effects on the transcriptome and non-linear correlation between copy number and expression (Ait Yahya-Graison *et al.* 2007; Huettel *et al.* 2008). In *Drosophila melanogaster,* buffering mechanisms were proposed to act through the gene regulatory networks to dampen the effect of gene dosage (Zhang *et al.* 2010; Malone *et al.* 2012). In *Saccharomyces cerevisiae,* gene expression largely correlated with copy number with some variation between genes (Gasch *et al.* 2016; Torres *et al.* 2016; Taggart and Li 2018).

While there is no effective compensation of chromosomal aneuploidies for the rest of the genome, X chromosomes are highly regulated in many animals. Typically, females (XX) contain twice the number of X chromosomes than males (XY) but the two sexes express similar levels of X chromosomal transcripts. Strategies that equalize X chromosome expression between sexes differ in different animals. In mammals (human and mouse) one of the X chromosomes is silenced in XX females, in flies (*Drosophila*) the single X chromosome is upregulated two-fold in XY males, and in worms (*Caenorhabditis*) both X chromosomes are downregulated by a factor of two in XX hermaphrodites (Meyer 2005; Samata and Akhtar 2018; Dossin and Heard 2021). While strategies differ, in each system a multi-subunit protein complex specifically targets and regulates X chromosome transcription in one of the two sexes through epigenetic mechanisms (Ercan 2015)

A high rate of copy number variations (CNV) was reported in mutation accumulation experiments in *Caenorhabditis elegans* (Lipinski *et al.* 2011; Konrad *et al.* 2018). In addition, 80% of the genome could be isolated as chromosomal duplications in the laboratory, leading to the conclusion that the worm is relatively tolerant to increased gene dosage (Hodgkin 2005). To understand the effect of increased gene dosage in *C. elegans,* we characterized large chromosomal duplications in several strains using DNA-seq and analyzed the effect of increased gene dosage in three chromosomal duplication and one fosmid insertion strain using mRNA-seq. Like other species, there was a complex response to increased gene dosage in *C. elegans.* While the average mRNA level increased for genes located within large duplications and the integrated fosmid, genes within the same chromosomal duplication showed varying levels of mRNA increase. An X chromosomal duplication that recruits the somatic dosage compensation complex showed lower mRNA increase compared to one that did not recruit, also demonstrating the contribution of epigenetic regulation to the effect of gene dosage.

## MATERIALS AND METHODS

### Strains

Unless otherwise noted, strains were maintained at 20°C on NGM agar plates using standard *C. elegans* growth methods. TY1916 (yDp11 (X;IV); lon-2(e678) unc-9(e101) X) contains duplication (yDp11), producing long, non-Unc homozygous hermaphrodites. SP117 (mnDp10 (X;I); unc-3(e151) X) produces wild type looking homozygous hermaphrodites. SP219 (mnDp1 (X;V)/+ V; unc-3(e151) X) contains the duplication (mnDp1), which is homozygous lethal, thus wild type looking heterozygous hermaphrodites containing the duplication were picked. BC4289 (sDp10 (IV;X)) contains the homozygous-viable duplication (sDP10). SP1981 (unc-115(mn481) dpy-6(e14) X; stDp2 (X;II)/+) contains homozygous-lethal duplication (stDP2). Wild type looking heterozygous hermaphrodites were picked. VC100(unc-112(r367) V; gkDf2 X) contains the homozygous viable deletion (gkDf2) and produces wild type looking hermaphrodites. OP37 (wgIs37 [pha-4::TY1::EGFP::3xFLAG + unc-119(+)]) is wild type looking and generated by (Sarov *et al.* 2012).

### DNA-seq

At least 20 worms were hand-picked as young adults. Worms were washed by settling animals at least 3 times with 1mL M9 and starved overnight to remove gut bacteria. Following a final M9 wash, worms were resuspended in 100uL TE and frozen. For DNA isolation, 400 μl of lysis buffer (0.1 M Tris-HCl; 0.1 M NaCl; 50 mM EDTA; 1.25% SDS) was added and worms were sonicated using Bioruptor 30 sec on/off at high for 30 min. Sonicated DNA was isolated using Qiagen MinElute kit and Illumina DNA sequencing libraries were prepared as described previously (Albritton *et al.* 2014). Single- or paired-end sequencing was performed using Illumina HiSeq-2000, and aligned to genome version WS220 (ce10) using Bowtie2 (version 2.3.2) with default settings (Langmead and Salzberg 2012). All replicate and read number information is provided in Supplemental File 1. Samtools version 1.6 (Li 2011) was used to merge replicates before running bamCompare from Deeptools version 3.3.1 (Ramirez *et al.* 2016), using the following options: -binSize 500, --scaleFactorsMethod None, --normalizeUsing CPM, --operation log2, -- minMappingQuality 30, --outFileFormat bedgraph, --ignoreDuplicates. Copy number analysis was performed with CNVnator version 0.3.3 (Abyzov *et al.* 2011) comparing data from mutant strains to reference genome WS220 (ce10) using bin_size = 1000. CNVnator output files listing deletions and duplications are provided in Supplemental File 2. Overlap of CNVs with genes were determined by Galaxy (https://usegalaxy.org/) tools using coverage option in “Operate on Genomic Intervals” (Afgan *et al.* 2018) and provided in Supplemental File 3. Only genes within the duplications and deletions greater than 10kb were considered for further analysis.

### mRNA-seq

Mixed stage embryos were collected from N2, TY1916 and SP117 by bleaching gravid adults. L2-L3 worms were collected by plating embryos and growing for 22-26 hours at 22.5°C. Worms were resuspended in at least 10 volumes of Trizol and stored at −80°C. RNA preparation was performed as previously (Albritton *et al.* 2014). Briefly, samples were freeze cracked three to five times, followed by TRIzol purification, and cleaned up with Qiagen RNeasy kit. From 0.5-10 μg of total RNA, mRNA was purified using Sera-Mag oligo(dT) beads (Thermo Scientific), sheared and stranded Illumina sequencing libraries were prepared using a previously published protocol (Parkhomchuk *et al.* 2009). Sequencing was performed with Illumina HiSeq-2000 and reads were aligned to genome version WS220 with Hisat2 version 2.2.1 (Kim *et al.* 2019) using default parameters. Count data was calculated using HTSeq version 0.13.5 (Putri *et al.* 2022) and differential expression was performed using the R package DESeq2 version 1.30.1 (Love *et al.* 2014). FPKM values were generated using Cufflinks version 2.2.1 with options -p 8 --library type fr- firststrand (Trapnell *et al.* 2010). FPKM and DEseq2 output are provided in Supplemental 4. In figure 3, log2 fold change values were median centered by subtracting the genome median from each value.

### Gene enrichment analysis in OP37

Genes differentially expressed in OP37 compared to N2 were analyzed using the Wormbase tool Gene Set Enrichment Analysis (Angeles-Albores *et al.* 2016). PHA-4 ChIP-seq binding peaks from OP37 L3 larvae were downloaded from the modERN project (Kudron *et al.* 2018). The results of the enrichment analyses and the list of differentially expressed genes are provided in Supplemental File 5.

### Data availability

A list of new and published data used in this study is provided in Supplemental File 1 with GEO accession numbers. The new data is available at GEO Series number GSE198682.

## RESULTS

### Characterization of chromosomal duplications and deletions using DNA-seq

To analyze the effect of gene dosage on mRNA expression, we used previously isolated strains with megabase scale duplications (Supplemental File 1). Since prior characterization of the duplications were done by visible genetic markers (Hodgkin 2005; Edgley *et al.* 2006), we performed DNA-seq to map genes that were duplicated or deleted. First, we calculated average read coverage within 1 kb windows tiled across the genome. Plotting the ratio of coverage to wild type confirmed previously mapped duplications (Figure 1).

**Figure 1.**
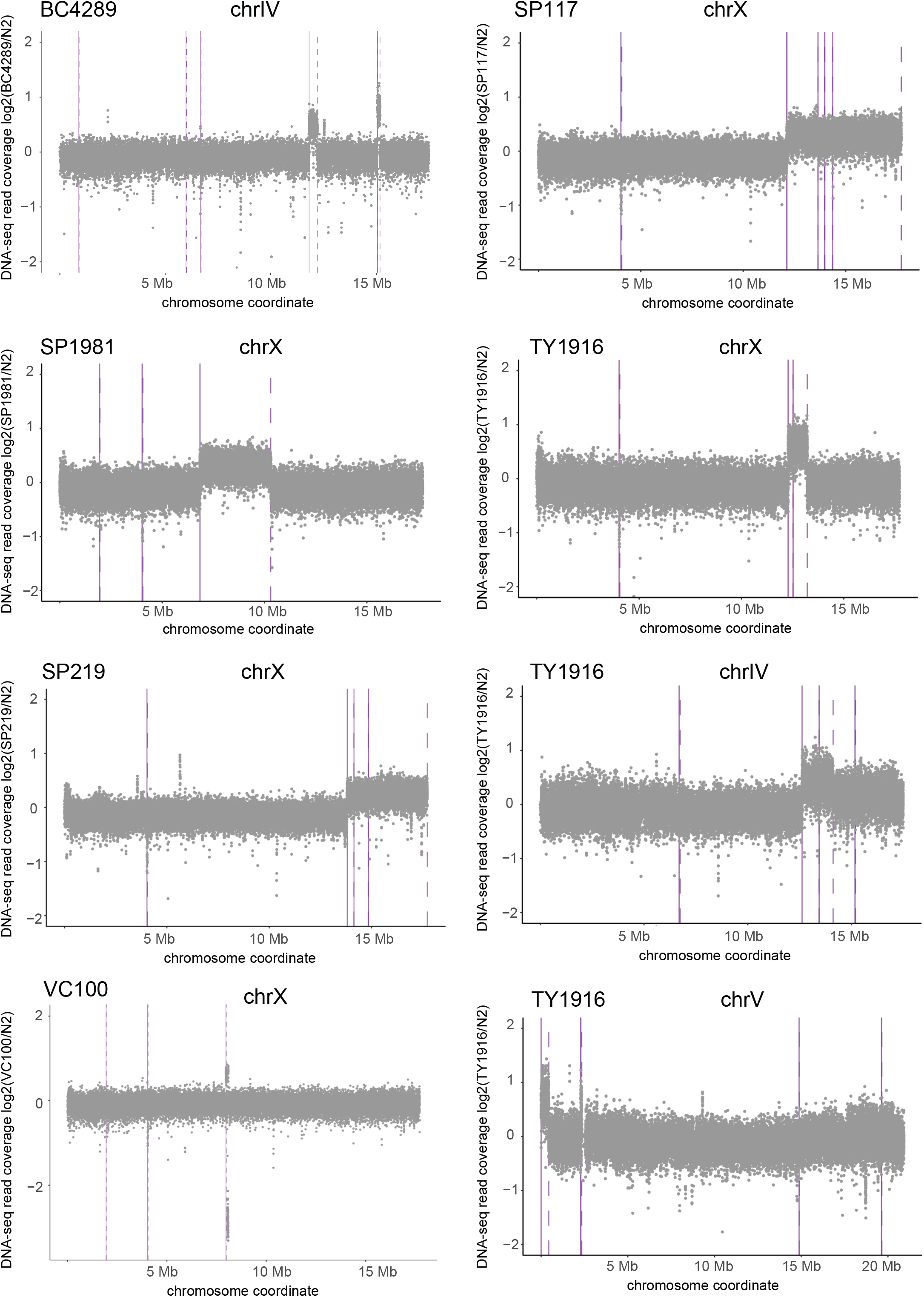
DNA-seq analysis of large chromosomal duplications. The x-axes are coordinates across the chromosomes containing duplications in each strain. The y-axis is log2(mutant/N2 control) coverage (CPM (counts per million) normalized and averaged for 1 kb windows). Solid vertical lines indicate the start of a duplication and dotted lines indicate the end of a duplication as determined by CNVnator output.

To further characterize the strains, we used CNVnator to generate a list of genomic windows with increased and decreased coverage compared to N2 (Abyzov *et al.* 2011) (Supplemental File 2). In addition to the mapped changes, many of the strains showed smaller CNVs that were different from the N2 strain (Figure 2). Some CNVs were common among the duplication strains, which may have originated from a laboratory strain polymorphic to N2 (Vergara *et al.* 2009). Notably, there were multiple smaller deletions and insertions within and at the boundaries of the larger duplications (Supplemental File 2). One example is the presence of a duplication near the ~40 kb deletion on the X chromosome in the VC100 strain. Presence of additional changes in the boundaries may be due to imperfect repair of the double strand breaks used to induce aneuploidies, followed by selection to laboratory conditions (Farslow *et al.* 2015).

**Figure 2.**
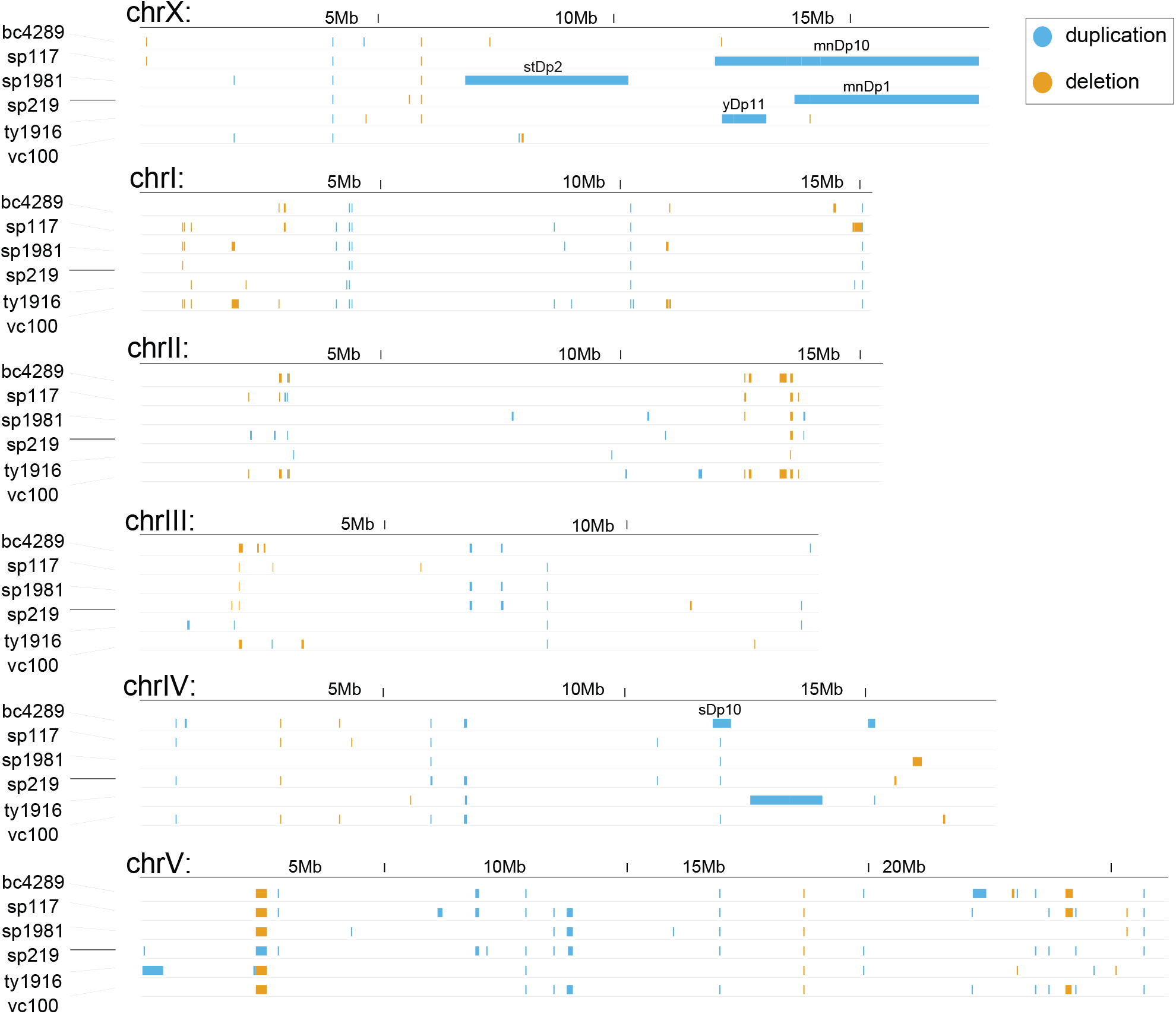
CNVnator-identified duplications and deletions. CNVnator identified duplications and deletions from each strain are visualized across the whole genome. Output files for each strain is provided in Supplemental File 2. Duplications are shown as blue, and deletions are shown as orange bars. The previously characterized large chromosomal duplications are labeled in each strain.

**Figure 3.**
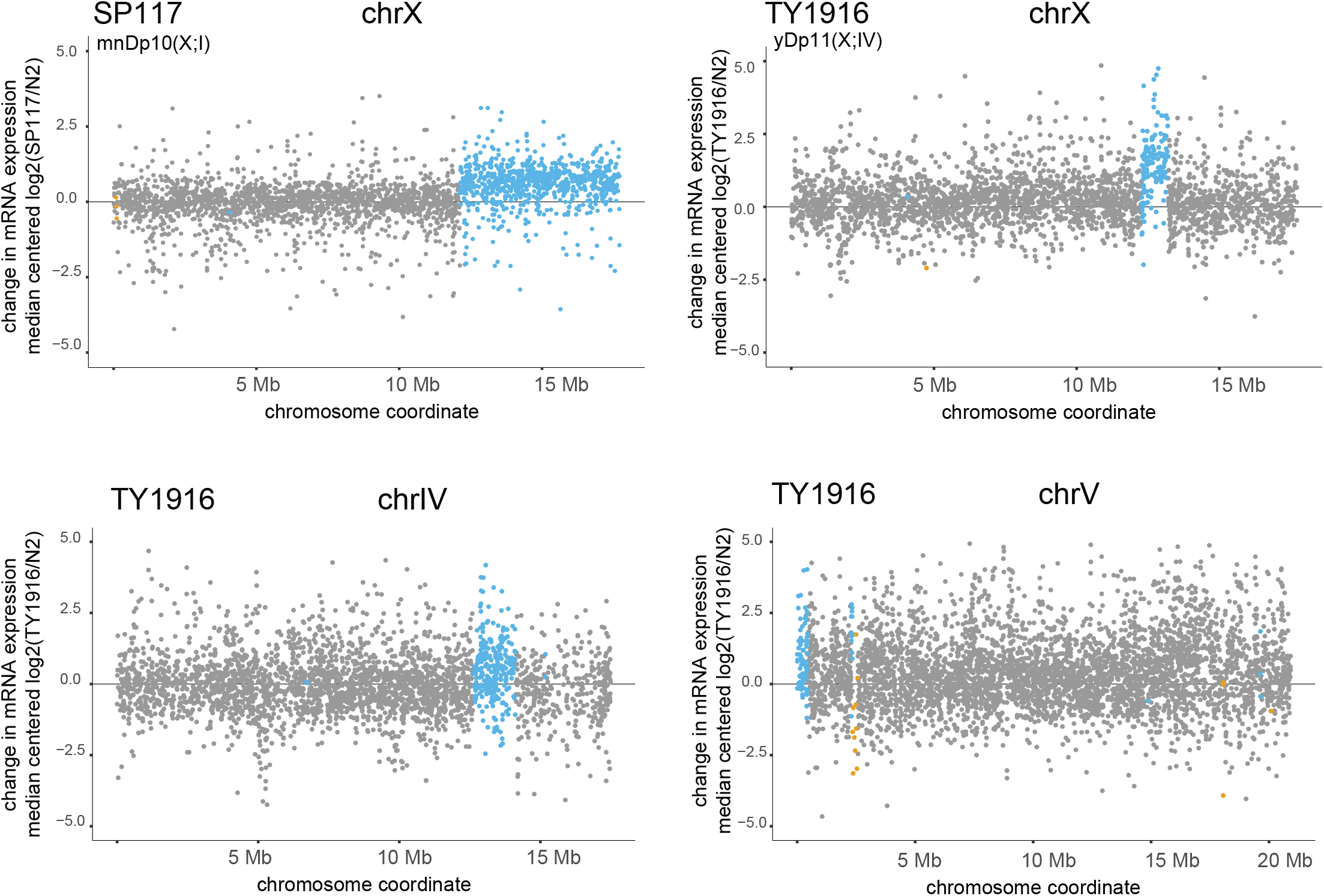
mRNA-seq analysis of four duplicated regions in two strains. Changes in mRNA expression in the SP117 and TY1916 strains compared to N2 are shown. In each graph, the x-axis shows the entire chromosomes from which the duplications originated. The y-axis is the genome-median centered log2 ratio (mutant/N2) as determined by DESeq. The DESeq outputs are provided in Supplemental File 4. The duplicated genes are highlighted in blue, deleted genes in orange, and unaffected genes in gray. Wilcoxon rank sum test p values are as follows: SP117 X-duplicated versus X-unaffected <2.2e-16, TY1916 X-duplicated versus X-unaffected <2.2e-16, TY1916 V-duplicated versus V-unaffected = 0.0016, TY1916 IV-duplicated versus IV-unaffected =0.0029. Gene categories for all analyzed strains are provided in Supplemental File 3.

### Increased gene dosage and mRNA expression from chromosomal duplications

To analyze the effect of chromosomal duplications on gene expression, we performed mRNA-seq in two homozygous viable strains with various length and number of duplications. SP117 contains an approximately 5 Mb X chromosomal duplication attached to chromosome I (mnDp10). TY1916 contains a ~1 Mb duplication from the X chromosome attached to chromosome IV (yDp11). TY1916 also contains previously uncharacterized changes involving the right arm of chromosome IV and the left end of chromosome V (Figure 1). The left tip of chromosome V coverage (0.67) is similar to that of the characterized duplication from the X chromosome (0.60), thus appears to be duplicated. In the case of region with increased coverage on chromosome IV, the median ratio of coverage is lower (0.40) than that of the X duplication (0.60), thus this region may be present as free duplication or heterozygous in the population of worms used for DNA-seq.

To identify the genes affected by the chromosomal duplications we used the CNVnator defined regions and categorized genes as duplicated or deleted (if transcription start-end of the gene fully overlaps with the CNV), affected (if overlap is 1 bp or more) or unaffected (no overlap) (Supplemental File 3). We then plotted the log2 ratio of mRNA coverage in the duplicated strain compared to control along the wild type chromosomes, highlighting genes within each category. Across all duplications, average mRNA-seq expression was increased, suggesting that gene dosage correlates with mRNA expression in *C. elegans* (Figure 3). Notably, the effect on individual genes was variable, showing a range of log2 ratios for genes located within the same duplication (Supplemental File 4).

In mRNA-seq analyses using ratios, inclusion of lowly expressed genes reduces the magnitude of observed effect (Deng *et al.* 2011). To address this problem, we filtered out genes with FPKM values less than 1 in any wild type replicate, and replotted the three large duplications in SP117 and TY1916 (Supplemental Figure 1A). The median mRNA-seq increase for all three duplications were similar with or without filtering (1.58 vs 1.51 for mnDp10, 2.34 vs 2.49 for yDp11, 1.51 vs 1.43 for duplication on chr IV in TY1916), thus lowly expressed genes did not significantly skew the analysis. We also plotted FPKM expression values for each gene between the large duplication strain SP117 and wild type (Supplemental Figure 1B). Although the correlation is noisier at lower FPKM values, the shift in mRNA level is clear for genes located at the duplication.

To probe further into variability, we addressed if the tighter scatter of log2 ratios on the X chromosome (Figure 3) is due to lower expression noise, which was shown for the single X chromosome upregulated by the dosage compensation complex in flies (Lee *et al.* 2018). To address noise, we calculated the coefficient of variation for each gene (filtering out those with FPKM<1) using wild type mRNA-seq data replicates in embryos, larvae, and young adults (Supplemental Figure 1C). Overall, there was less variation in larvae, likely due to temporal dynamics of embryogenesis in embryos and germ cell development in young adults adding variability between collection of worms for mRNA-seq replicates. In all three developmental stages, the median coefficient of variation for X chromosomal genes was in between other chromosomes, suggesting that dosage compensation in worms does not reduce expression noise below that of autosomes.

### Increased mRNA expression and indirect effects of a multi-copy integrated fosmid

While megabase-scale chromosome duplications showed an average increase in mRNA level, we wondered if smaller chromosomal segments with higher copy numbers also increase mRNA expression. To this end, we used a multi-copy fosmid integration strain, where GFP-3xflag tag was inserted to the C terminus of *pha-4* gene within a fosmid containing three other genes (Sarov *et al.* 2012). The fosmid was then integrated into the genome randomly and a transgenic line that was vigorous and expressing the GFP-tagged *pha-4* was selected. This strain named OP37 was used to study the binding sites of *pha-4,* a transcription factor required for several developmental processes including the pharynx (Gaudet and Mango 2002; Zhong *et al.* 2010). The copy number for the fosmid was calculated to be 5.6 (Sarov *et al.* 2012).

The genes within and surrounding where the fosmid originated from are shown along with the DNA read coverage highlighting the increased gene dosage (Figure 4A). The insertion site of the fosmid has not been mapped. Notably, the mRNA level of all four genes including *pha-4* was increased in the OP37 strain compared to N2 control (Figure 4B). The increase for *pha-4* was about ~2.4-fold suggesting either feedback repression or an effect of GFP tagging on mRNA expression or detection. The mRNA level of the other three genes on the fosmid increased ~3.6-4.8 fold, reflecting the increase in their copy number.

**Figure 4.**
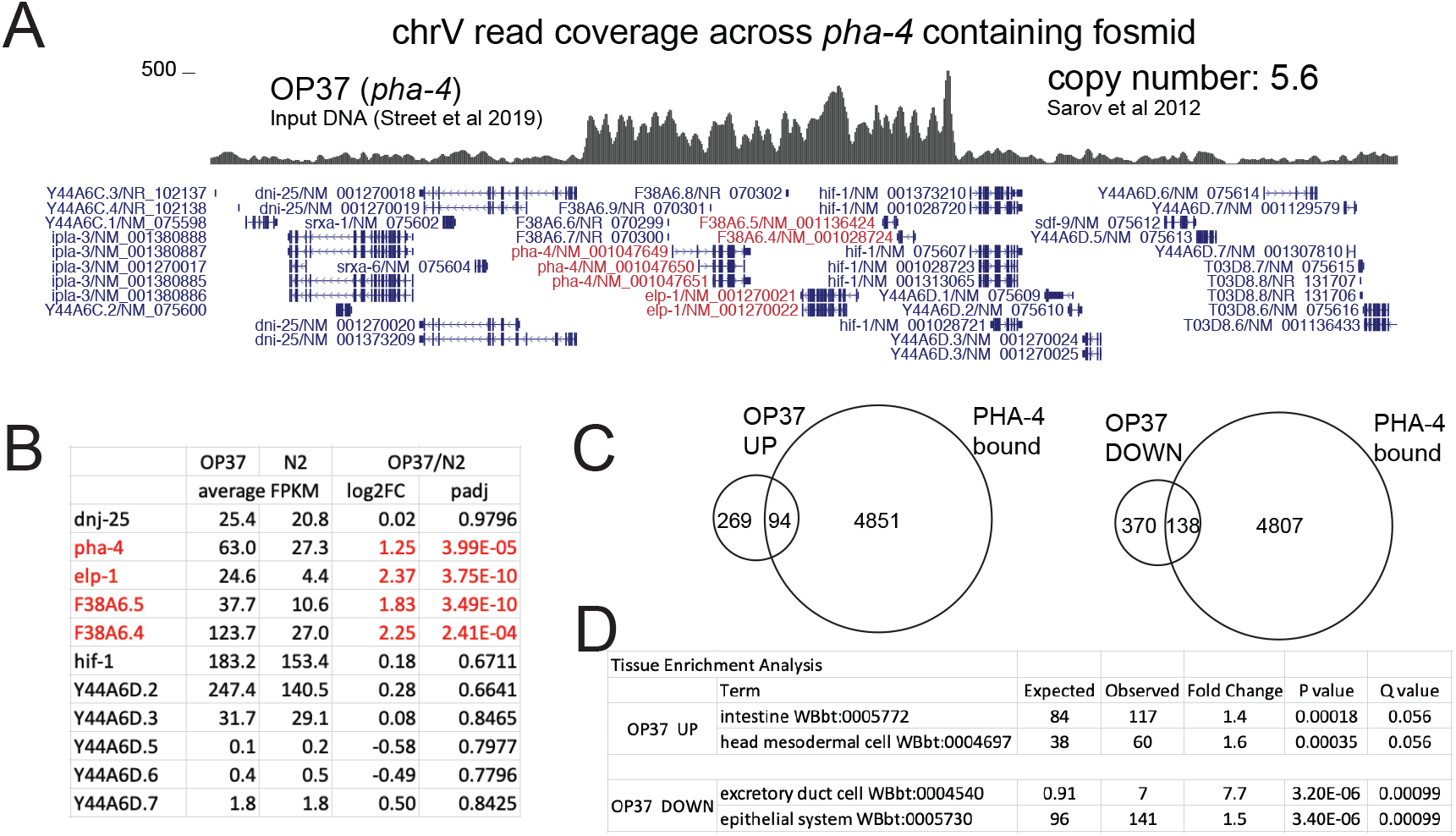
mRNA-seq analysis in a multi-copy integrated fosmid strain. A) DNA-seq read coverage plot of OP37 visualized on UCSC genome browser, centered at a ~70kb window across the region where *pha-4* containing fosmid originated from. Genes included in the fosmid are highlighted in red. B) Table showing the average FPKM, log2foldchange, and padj values from DESeq analyses, listed for genes within and neighboring the *pha-4* containing fosmid. The genes within the fosmid are highlighted in red. The entire analysis is included in Supplemental File 4. C) Overlap of genes bound by PHA-4 (modENCODE ChIP-seq peaks at 1 kb promoters) and genes up or downregulated in the OP37 strain compared to N2. D) Tissue enrichment analysis of differentially expressed genes in the OP37 strain. Top two categories are shown for up and downregulated genes, and the rest are provided in Supplemental File 5.

Highlighting the indirect effect on the transcriptome, 871 genes were identified to be up or downregulated in the OP37 strain, despite the fosmid including only four genes. Analysis of genes that were differentially expressed revealed no enrichment for PHA-4 bound genes (25% of upregulated and 27% of downregulated genes were bound by PHA-4 compared to 27% of all genes) (Figure 4C). It is possible that the ~2.4 fold increase in *pha-4* mRNA level does not cause an increase at the protein level or that the presence of multiple genes within the fosmid reduces *pha-4* specific changes on the transcriptome. The differentially expressed genes in the OP37 strain showed tissue specific enrichment possibly reflecting the developmental functions of *pha-4* and the neighboring genes (Figure 4D) (Supplemental File 5).

### The mRNA increase from an X duplication differs across developmental stages

Next, we wondered how epigenetic regulation of the X chromosomes in different tissues would affect the consequence of an X chromosomal duplication. To address this question, we first considered the germline, where the X chromosomal genes are repressed compared to autosomal genes during early meiosis (Schaner and Kelly 2006). We reasoned that if an X chromosomal duplication is also subject to repression, the mRNA increase would be lower in adult worms. Indeed, genes located within the 5Mb X chromosomal duplication in the SP117 strain showed less increase in mRNA expression in young adults compared to embryo and larvae (Figure 5A). X chromosome repression in the germ cells is measurable in whole animals because germ cells outnumber the somatic cells in adults, leading to lower X chromosome expression compared to autosomes (Figure 5B). Thus, our results suggest that repression of genes within the 5Mb X chromosomal duplication in the germline negates the mRNA effect of the duplications specifically in this tissue.

**Figure 5.**
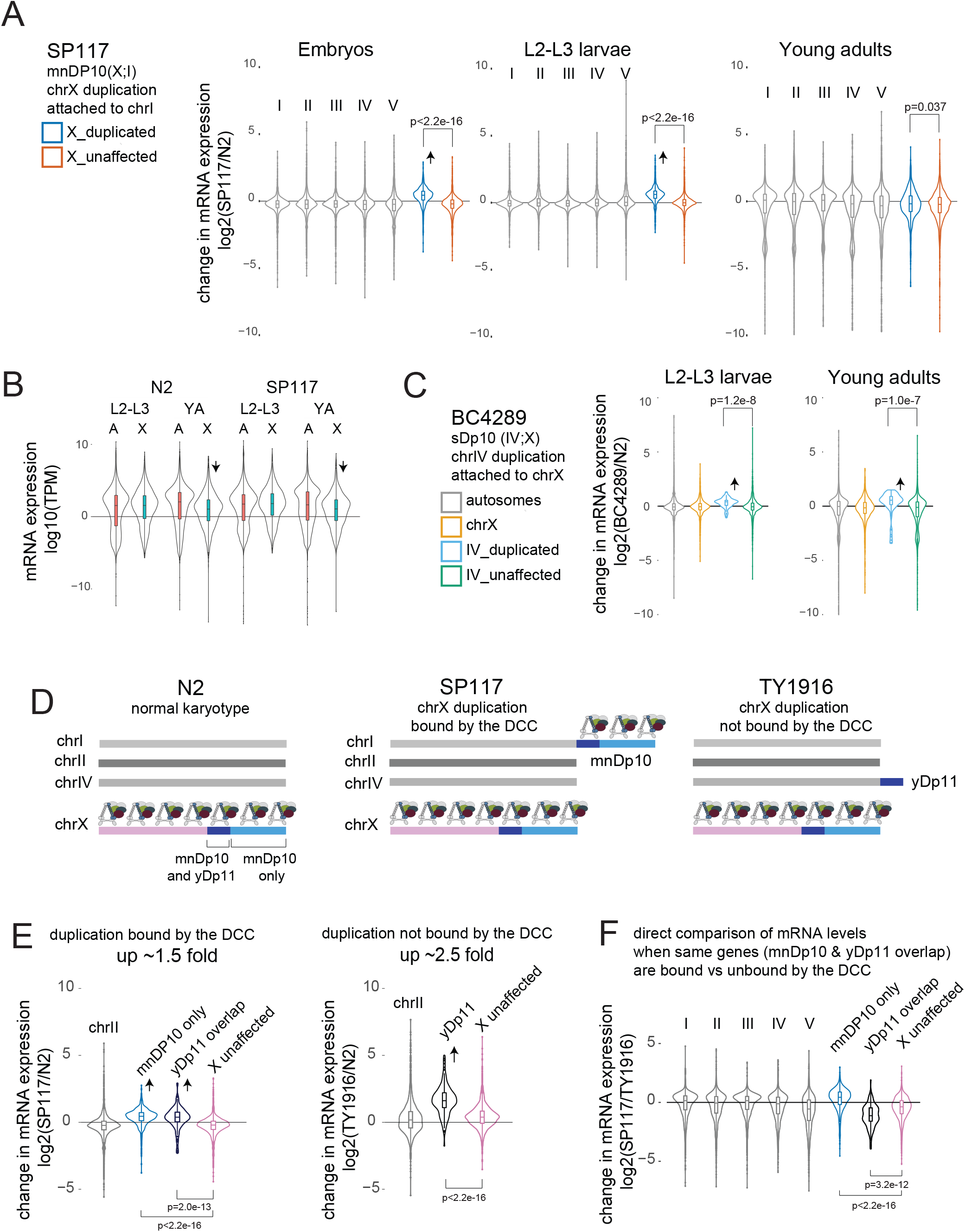
mRNA expression in X duplications subject to repression in the soma and germline. A) SP117 contains a large ~5 Mb duplication of chr X that is attached to chr I. Violin plots of log2foldchange from DESeq analysis of SP117 compared to N2 in embryos, L2-L3 larvae and young adult worms. Significance of difference between average log2 fold change for genes within and outside the duplication from chr X is tested by t-test. Increased mRNA expression from the duplication in embryos and larvae are highlighted with an arrow. B) Violin plot of mRNA levels (log10FPKM) from autosomes and the X chromosomes in larvae and in young adults in N2 and SP117. X chromosomes are repressed in the germ cells that are present in young adults but not in L2-L3 larvae. (One sided t-test (for repression) comparing average X chromosomal expression in L2-L3 larvae vs young adult (YA), p value N2=0.09, SP117=0.0005). C) BC4289 contains a ~1 Mb chr IV duplication attached to chr X. Violin plots of log2foldchange from DESeq2 analysis of BC4289 compared to N2 in L2-L3 larvae and young adults is shown. Significance of difference between average log2 fold change of genes within and outside the duplication from chr IV is tested using t-test. D) A schematic representing N2 control, SP117 and TY1916 genomes. The 1 Mb duplicated region (dark blue) in TY1916 (yDp11) does not bind to the somatic dosage compensation complex. The 4 Mb uniquely duplicated region in SP117 (mnDp10) is indicated in light blue. Within SP117(mnDp10) genes within the duplicated region that overlap with yDp11 (dark blue) is bound by the dosage compensation complex. E) Violin plots of log2foldchange of SP117 and TY1916 compared to N2 in L2-L3 larvae. 1.5 and 2.5 fold refer to the ratio of the median for the 1 Mb common duplicated region compared to the unaffected X chromosomal genes (lowly expressed genes are filtered (FPKM<1)). The arrows indicate the mRNA-seq increase in the duplicated regions, with p values generated from t-test. F) Violin plot of log2foldchange ratio for SP117 directly compared to TY1916. The increase in mRNA expression of the same set of genes is lower in SP117 when the ~1 Mb X chromosome duplication is bound by the DCC compared to when unbound in TY1916.

The duplication strain BC4289 contains a ~1 Mb translocation from chromosome IV to X allowing us to test if autosomal genes attached to the X chromosome are repressed in germ cells. Unlike the 5 Mb X-to-I translocation in SP117, the 1 Mb IV-to-X translocation in BC4289 showed a similar increase in mRNA expression in young adults compared to larvae (Figure 5C). Thus, germline repression may not affect autosomal genes even when they are physically attached to the X chromosome. In the future, multiple autosome to X duplications should be analyzed to test the generality of this conclusion. Nevertheless, the SP117 data suggest that epigenetic regulation of X chromosomal genes reduces the effect of a large X-duplication on mRNA expression in the germline.

### Somatic dosage compensation reduces the level of mRNA increase in X chromosomal duplications

In *C. elegans,* hermaphrodite X chromosomes are repressed by a factor of two by somatic dosage compensation (Kramer *et al.* 2016). This repression is mediated by a condensin-based dosage compensation complex (DCC) that specifically binds to and represses transcription initiation from both X chromosomes (Kruesi *et al.* 2013; Kramer *et al.* 2015). DCC is recruited specifically to the X chromosomes by several cis-regulatory elements on the X (McDonel *et al.* 2006; Ercan *et al.* 2007; Jans *et al.* 2009; Albritton and Ercan 2017). Recruitment leads to spreading of the complex to nearby chromatin, creating robust binding across the X chromosomes (Albritton *et al.* 2017; Street *et al.* 2019).

Previous studies used immunofluorescence to analyze the ability of several X chromosomal duplications to recruit the DCC (Csankovszki *et al.* 2004; Blauwkamp and Csankovszki 2009). These studies found that the ~1 Mb duplication in TY1916 (yDp11) does not recruit the DCC. Although one strong and one weak recruitment site is present within yDp11, insertion of few recruitment elements is not sufficient to recruit high levels of DCC to an autosome (Albritton *et al.* 2017), explaining the lack of robust DCC binding to this shorter duplication. The same stretch of DNA within the larger~5 Mb duplication in SP117 (mnDP10), is bound by the DCC (Csankovszki *et al.* 2004), due to the additional ~4 Mb region containing five strong and fifteen weaker recruitment elements. Since genes in yDp11 are also in mnDP10, we were able to test if epigenetic repression by the DCC reduced the effect of increased dosage across the shared genes (Figure 5D). Indeed, the average mRNA increase from the 1 Mb region differed in the two strains (Figure 5E). The median increase of mRNA expression from the 1 Mb duplication was lower when bound by the DCC (SP117) compared to when unbound (TY1916).

In TY1916, there are four copies of genes located within the 1 Mb yDp11; two X chromosomal copies under DCC-mediated repression and two duplicated copies on chromosome IV without DCC. In SP117, all four copies of the genes within the commonly duplicated region between mnDp10 and yDP11 are under DCC-mediated repression. In both strains, the expected level of median mRNA increase (based on copy number and the assumed 2-fold repression by the DCC) was lower than that of observed; ~2.3 fold versus 3 in TY1916, and ~1.5 fold versus 2 in SP117 (after filtering genes (FPKM<1) median increase was ~2.5 for TY1916 and 1.5 for SP117). It would be interesting to know if other mechanisms of repression such as lamina-mediated organization of repressive chromatin contribute to repression of X chromosomal duplications (Snyder *et al.* 2016; Lee and Oliver 2018). Regardless, the lower level of mRNA increase from the same set of duplicated genes in the SP117 (DCC-bound) strain compared to TY1916 (DCC-unbound) (Figure 5F) indicates that DCC mediated repression reduced the consequence of increased gene dosage on mRNA expression.

## DISCUSSION

In *C. elegans,* several observations supported the idea that this organism is robust to changes in gene dosage in laboratory (Hodgkin 2005; Lipinski *et al.* 2011; Sarov *et al.* 2012; Konrad *et al.* 2018). Here we showed that tolerance to increased gene dosage is not due to a genome-wide compensation mechanism acting at the mRNA level. However, it remains possible that effective mechanisms of compensation exist at the protein level (Dalley *et al.* 1993; Chang and Liao 2020). Post-transcriptional control of gene expression is common in *C. elegans* particularly in the germline (Nousch and Eckmann 2013), a tissue whose function is central to the viability of all isolated strains.

Our analysis of chromosomal duplications in *C. elegans* indicate that similar to other organisms (Kojima and Cimini 2019), increased gene dosage leads to increased mRNA expression with a high degree of variability between genes (Figure 3). The variation in response to gene dosage could be due to measurement errors based on technical and biological variation in mRNA expression. It could also be due to feedback through gene regulatory networks (Malone *et al.* 2012). Indirect effect of increased gene dosage on the transcriptome was particularly evident in the fosmid integration strain (Figure 4). It is possible that compensatory changes in the expression of other genes render chromosomal duplications and copy number variations viable in the laboratory.

Partial aneuploidies and copy number variations in specific genes have been implicated in a wide range of diseases (Inaki and Liu 2012; Tang and Amon 2013; Durrbaum and Storchova 2016). To determine the cause and consequences of these copy number variations on organism phenotypes, it is necessary to understand the variation both between genes and between tissues. Here, our results demonstrate that tissue-specific mechanisms of X chromosomal gene repression control the level of mRNA increase from certain duplications in the germline and the soma. Thus, increased gene dosage could result in varying levels of mRNA-seq increase, when the same set of genes are under different epigenetic regulation. Overall, our work adds *C. elegans* to the line of research analyzing partial aneuploidies, demonstrating a lack of genome-wide dosage compensation mechanism acting at the mRNA level, highlighting gene-to-gene variability and the contribution of epigenetic regulation to the mRNA response to gene dosage.

---

**Supplemental Figure 1.**
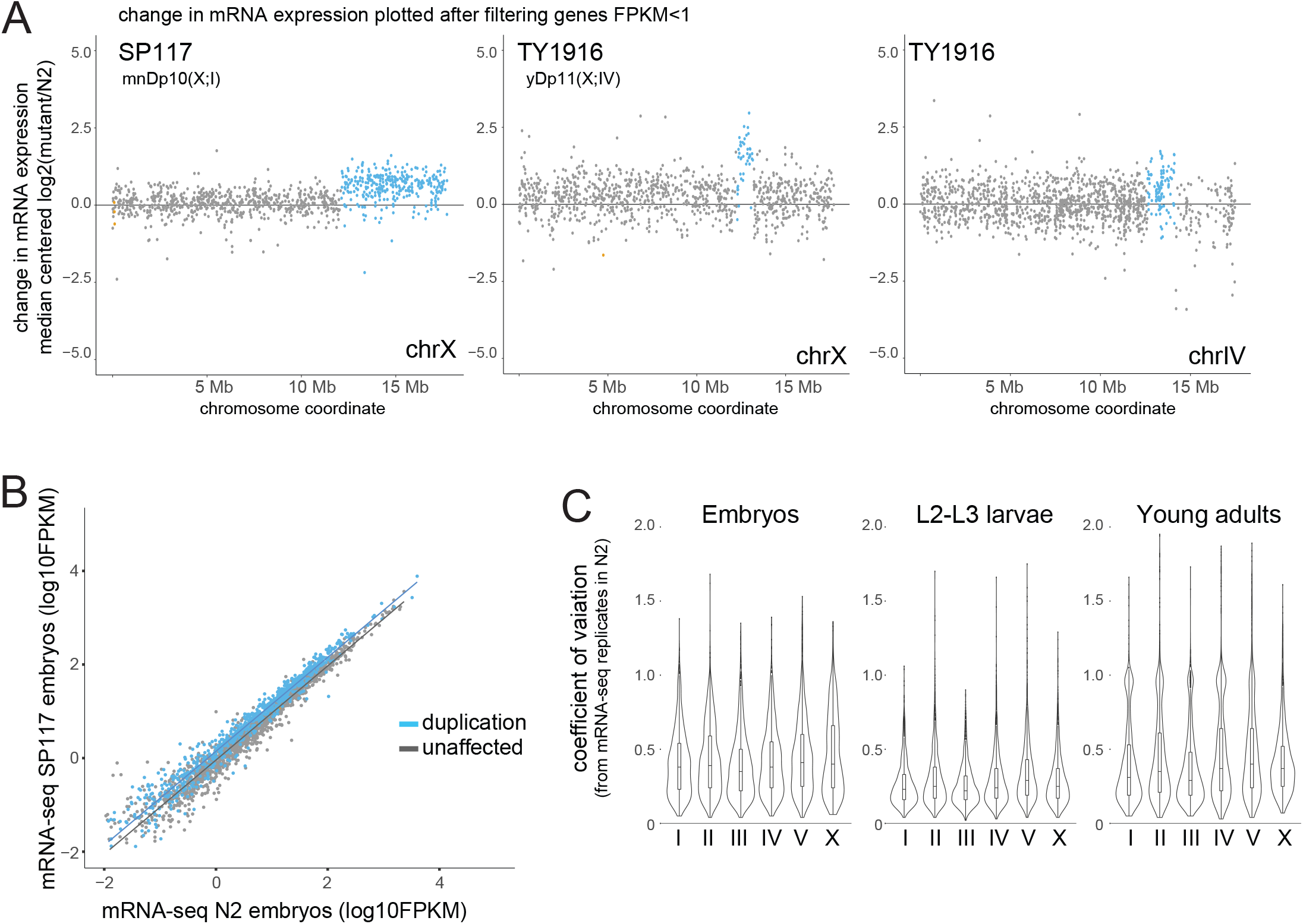
A) Changes in mRNA expression in the SP117 and TY1916 strains compared to N2 are shown. In each graph, the x-axis shows the entire chromosomes from which the duplications originated from. The y-axis is the genome-median centered log2 ratio (mutant/N2) as determined by DESeq2. Genes that are lowly expressed (FPKM<1 in any wild type mRNA-seq replicate) have been removed. The duplicated genes are highlighted in blue, deleted genes in orange, and unaffected genes in gray. B) Scatter plot analysis of FPKM expression level for the X chromosomal genes in the SP117 strain. The duplicated genes are shown in blue. Trendlines were fitted in R using ggplot’s linear model (lm()) function. D) Expression noise for each chromosome is analyzed through calculating the coefficient of variation from FPKM values of mRNA-seq replicates. N2 wild type data was used and genes that showed FPKM<1 in any of the replicates were removed.

## ACKNOWLEDGEMENTS

Research reported in this publication was supported by NIGMS of the National Institutes of Health under award number R01GM107293 and R35GM130311. We thank Marissa Knoll for contributing to the OP37 strain collection. Some strains were provided by the CGC, which is funded by NIH Office of Research Infrastructure Programs (P40 OD010440).

## Notes

### Competing Interest Statement

The authors have declared no competing interest.

### Summary of Updates

The Manuscript now includes a supplemental file 1, where we have analyzed the impact of lowly expressed genes on the gene expression ratios.

